# Dual Role of LH-GABA Neurons in Encoding Alcohol Reward and Aversive Memories

**DOI:** 10.1101/2025.10.22.683984

**Authors:** Isis Alonso-Lozares, Dustin Schetters, Yvar van Mourik, Allison J. McDonald, Taco J. De Vries, Nathan J. Marchant

**Author notes:** Corresponding Author: Nathan Marchant.

## Abstract

One of the core aspects of alcohol use disorder is continued use despite negative consequences. Individuals with an alcohol use disorder typically engage in behaviors which represent a failure to integrate the aversive consequences of their actions. The neural mechanisms of learning about conflicting experiences remain poorly understood. Previous research has highlighted the critical role of GABAergic neurons in the Lateral Hypothalamus (LH-GABA) in rewarding memories. However, whether this role extends to aversive outcomes, and competition between rewarding and aversive outcomes remains unexplored. In this study, we sought to elucidate the role of LH-GABA neurons in encoding and expressing alcohol reward and aversive memories using fiber photometry calcium imaging. We used a dual-virus approach to confine expression of jGCaMP7f to GABAergic neurons in LH of male and female Long-Evans rats. In our first experiment, following acquisition of a cue-alcohol association, a mild foot shock was introduced on 50% of the trials. In the second experiment, we used three different cues to signal alcohol, foot-shock, or no consequence. Subsequently, we combined these conditioned stimuli in pairs to evoke motivational conflict. Our results reveal that LH-GABA activity is associated with cues predictive of both appetitive and aversive outcomes and is highest to a cue predictive of an aversive outcome. Additionally, the response of LH-GABA activity to an aversive shock stimulus is attenuated in the presence of alcohol. In summary, our findings show the activity of LH-GABA neurons is involved in learning both appetitive and aversive associations, and their interaction in conflict.

**Significance statement.:** This study addresses a critical knowledge gap in understanding the neural mechanisms behind learning in conflicting situations. Focusing on the GABAergic neurons in Lateral Hypothalamus, which have previously been associated with rewarding memories and motivation, we explore their role in the competition between rewarding and aversive motivation. We used fiber photometry to record LH-GABA neurons and found that their activity is heightened for aversive outcomes, irrespective of certainty. Notably, in situations of motivational conflict, this activity to the shock is dampened in the presence of alcohol. These insights not only enhance our understanding of the neural basis of emotional adaptation but also hold implications for psychiatric disorders like alcohol use disorder, depression, and anxiety, providing potential avenues for therapeutic interventions.

## Introduction

Our actions are often guided by the anticipation of rewards or the avoidance of aversive outcomes, and we develop expectations for these outcomes through learning about environmental stimuli that reliably predict these outcomes. Previous research has extensively examined the neural mechanisms underlying the learning of reward and aversive predictive stimuli in isolation (LeDoux, 2000; Cardinal et al., 2002). How these systems interact has been the focus of past behavioural work (Konorski, 1967; Dickinson and Dearing, 1979; Peck and Bouton, 1990), and recent work has investigated the neural substrates of competition between these motivational states (Burgos-Robles et al., 2017; Illescas-Huerta et al., 2021; Choi et al., 2022). In complex environments, adaptive decision-making is crucial for survival (Miller, 1971). In humans, decision-making under conflict is essential for promoting psychological wellbeing. Maladaptive responses to conflict can contribute to psychological disorders such as anxiety (Gray and McNaughton, 2000), depression (Dickson and MacLeod, 2004; Yechiam et al., 2008; Chandler et al., 2009), obsessive compulsive disorder (Figee et al., 2011; Milad et al., 2013), and substance use disorder (Barkby et al., 2012; Naqvi et al., 2014; Luscher et al., 2020).

Approximately 80% of reported cases of drug abuse are alcohol-related (Heinz et al., 2009), and alcohol use disorder has been characterized by continuous alcohol use despite increasing negative consequences (Barkby et al., 2012), and a high likelihood of relapse (Hunt et al., 1971). In alcohol addiction, initially strong positive alcohol associations create anticipation of positive outcomes and reinforce an individual’s motivation to seek them (O’Brien et al., 1992). The persistence of positive associations is understood to be a critical component of the high likelihood of relapse (Bossert et al., 2013; Sliedrecht et al., 2019; Suzuki et al., 2020). During the development of alcohol use disorder, the consequences of alcohol use become negative, yet alcohol use persists (Hasin et al., 2013; Hopf and Lesscher, 2014). The specific neural substrates and behavioral dynamics underlying this shift from positive reinforcement to negative consequences remain poorly understood. Unraveling these intricacies could significantly advance our comprehension of addiction mechanisms and inform targeted interventions for individuals with alcohol use disorder.

Recently, we and others have shown that the GABAergic subpopulation of neurons in Lateral Hypothalamus (LH-GABA) is involved in reward and aversive learning processes (Sharpe et al., 2017; Sharpe et al., 2021; Alonso-Lozares et al., 2024). This subpopulation is functionally and anatomically distinct from Orexin (Ox) and Melanin-Concentrating Hormone (MCH) populations (Karnani et al., 2013; Jennings et al., 2015). LH-GABA have reciprocal projections with Ventral Tegmental Area (VTA) (Nieh et al., 2016), Nucleus Accumbens Shell (NAcSh) (O’Connor et al., 2015), and Lateral Habenula (LHb) (Lazaridis et al., 2019; Trusel et al., 2019), which are areas also involved in both reward and aversive learning and motivation. We previously found that LH-GABA activity is associated with encoding, extinction, and reinstatement of alcohol-predictive cues (Alonso-Lozares et al., 2024), and opto-inhibition of LH-GABA activity prevents learning of cue-food, alcohol, and shock associations (Sharpe et al., 2017; Sharpe et al., 2021; Alonso-Lozares et al., 2024)

Here, we sought to investigate the role LH-GABA neurons play in the integration of reward and aversive learning using fiber photometry calcium imaging (Nakai et al., 2001; Cui et al., 2014). We first showed that LH-GABA activity significantly increases to a cue after the consequences change from alcohol alone to alcohol with a potential shock. In the second experiment, we trained animals to associate three different cues with certain alcohol, shock, or no consequence, and in the next phase, we presented the cues in pairs to simulate motivational conflict. We found that LH-GABA activity is consistently highest for the cue which predicts shock compared to alcohol, whether presented alone or in compound with the alcohol predictive cue. We also showed that the activity of LH-GABA neurons to the shock is decreased when this occurs in combination with alcohol reward. These data suggest that the contribution of LH-GABA extends beyond reward-only associative learning and might also play a role in aversive learning.

## Materials and methods

### Subjects

We used 19 adult wild-type Long-Evans rats (3 males and 16 females), from Janvier (France). All behavioural and surgical procedures were performed in the animal facility of the Vrije Universiteit, Amsterdam, The Netherlands. Rats were kept in temperature and humidity-controlled rooms, in an inverted 12/12 light/dark cycle, and we trained the rats during the dark cycle. Food and water were provided ad libitum. Animals were kept in cages of either 2 or 3 cage mates. All procedures were approved by the Instantie voor Dierenwelzijn of the Vrije Universiteit, Amsterdam, The Netherlands (CCD: AVD1140020187084).

We did not make specific analysis to determine sample size prior to any experiments. Cohorts were balanced by experimental group prior to the experiment starting. We did not exclude any rats based on behavioural reasons. Rats that did not show expression and fiber placement within LH were excluded from analysis. Experiment 1 consists of 13 rats (13F) and Experiment 2 consists of 6 rats (3F/3M).

### Apparatus

All procedures were performed in standard Med Associates operant chambers with data collected through the MED-PC V program (Med Associates, Georgia, VT, USA). Each chamber had a grid floor and a large opening at the top to allow patch cords through. Each chamber was equipped with a white noise generator, and one or two light cues, as well as a magazine port where the alcohol (20% ethanol) was infused into the receptacle via tubing connected to a 20ml syringe controlled by a Razel pump. The magazines were equipped with a beambreak device to measure total time spent in the magazine. The grid floor consisted of 19 rods (VFC-005) and was connected to the Med Associates Aversive Stimulator/Scrambler. Custom made Med-PC programs controlled all aspects of the task.

For fiber photometry, excitation and emission light was relayed to and from the animal via optical fiber patch cords (0.48 NA, 400-µm flat tip; Doric Lenses). Blue excitation light (490 or 470 nm LED [M490F2 or M470F2, Thorlabs]) was modulated at 211 Hz and passed through a 460–490 nm filter (Doric Lenses), while isosbestic light (405 nm LED [M405F1, Thorlabs]) was modulated at 531 Hz and passed through a filter cube (Doric Lenses). GCaMP7f fluorescence was passed through a 500- to 550-nm emission filter (Doric Lenses) onto the photoreceiver (Newport 2151). Light intensity at the tip of the fiber was measured before every training session and kept at 21 µW. A real-time processor (RZ5P, Tucker Davis Technologies) controlled excitation lights, demodulated fluorescence signals and received timestamps of behavioural events. Data were saved at 1017.25 Hz and analysed with custom-made Matlab scripts, available at: https://github.com/ialozares/AlcoholConditioning.

### Drugs

Alcohol solutions were prepared using 70% (v/v) ethanol diluted with autoclaved tap water and were administered using normal bottles or syringe pumps in operant chambers.

### Surgeries

The day before and 30-60 minutes prior to surgery, we injected rats with the analgesic Rymadil® (5 mg/kg; Merial, Velserbroek, The Netherlands) Surgery was performed under isoflurane gas anesthesia (4% induction, 2% maintenance, PCH; Haarlem). The surgical area on the skull was shaved & disinfected before surgery and the skin was locally anesthetized using a subcutaneous injection of 1.0 ml/kg 10% lidocaine.

Craniotomies above were performed, followed by AAV injections using a micro-syringe connected to a micropump (UltraMicroPump 4, WPI surgical instruments), which was lowered into the brain until it reached the DV coordinates (see below for details). 6 screws were inserted into the skull to support the cement needed for fiber anchoring. Next, an optic fiber was implanted using the same stereotactic apparatus and bregma-relative coordinates as used for the AAV injection, with the fiber tip 0.5 mm above the injection depth. The optic fiber implant(s) (when applicable) were then secured to the skull using dental cement (IV Tetric EvoFlow 2g A1, Henry Schein, Almere). Rymadil (5 mg/kg; s.c.) was administered for 2 days after the surgery. Rats were given one week of recovery following surgery.

### Adeno-associated Viruses (AAV)

For both experiments, we injected a cocktail of one AAV expressing Cre under the GAD promotor (ssAAV-5/2-hGAD67-chI-iCre-SV40p(A)/ ssAAV-9/2-hGAD67-chI-iCre-SV40p(A), UZH-vector core, titer: 5.6x10^12) and another AAV encoding Cre-dependent jGCamP7f (ssAAV-5/2-hSyn1-chI-dlox-jGCaMP7f(rev)-dlox-WPRE-SV40p(A), UZH-vector core, titer: 2.4x10^12). 1µl of AAV solution was injected unilaterally into LH (AP: + 2.3, ML: 2, DV: -8.4 and –8) over two infusions. Each infusion was 2.5 min duration, and the needle was left in place for 5 min. We first infused 0.5 µl at DV: -8.4mm and then after infusion and diffusion we injected 0.5 µl at DV: -8.0mm. A 400µm diameter optic fiber (Doric Lenses) was implanted above LH.

### Behavioural procedures

#### Intermittent-access 2-bottle-choice alcohol access in the homecage

For all behavioural experiments, we first habituated rats to alcohol consumption using an alcohol homecage procedure (Wise, 1973; Simms et al., 2008). Rats had intermittent 24h access to a bottle containing 20% ethanol (EtOH), and a bottle containing water on Monday, Wednesday, and Friday. After 24h, or 48h over the weekend, the bottles were weighed and swapped for a single water bottle for the following 24h. These two phases were repeated 12 times. Additionally, we used two control empty cages without animals to account for spillage. The overall consumption of alcohol per rat after each session was measured in grams, by subtracting pre- and post-measurements for each cage and the average of the two control cages, accounting for 2-gram spillage and multiplying by the density of 20% ethanol (0.97).

During this phase, we habituated the rats to the photometry patch cords by tethering them to the patchcord and leaving them in their designated behavioural boxes for 30 minutes on the alcohol off days (Tuesday and Thursday). Alcohol magazines were removed from the boxes during habituation.

#### Pavlovian conditioning

Rats were taken from their home-cage housing room and moved to the training room containing the chambers and the photometry recording setup. We placed the rats in the chambers and trained them once a day during weekdays. Sessions lasted from 20 – 45 minutes depending on the experiment and the phase. We measured the time spent in the magazine, which was indicated by the rat breaking the beam within the magazine throughout the entire cue period.

### Experimental design

#### Exp. 1: Monitoring LH-GABA neurons during reacquisition of a cue-alcohol association, and during appetitive-aversive conditioning

In this experiment, prior to the behavioral tests described below, we trained rats to associate a cue with alcohol reward, extinguished that cue, and then tested for reinstatement. Figure S1 shows the experimental procedure of these phases, and the behavioral data associated with it. These data are presented in detail in Alonso-Lozares et al, (2024). Following this, we re-trained the rats on the original alcohol Pavlovian conditioning task, described below. The conditions of the original training phase are identical to the ‘Reacquisition’ Phase described below.

##### Reacquisition Phase

We retrained the rats on the previously learned association between a cue with alcohol delivery in the alcohol magazine (CS+ trials) and a different cue with no programmed consequences (CS-trials). Cues consisted of two different stimuli (white noise or houselight), counterbalanced across rats. In each session, rats received 8 presentations of each CS (16 trials total). Data were further divided into three phases (pre-CS, during-CS, and post-CS), each lasting 10 seconds and resulting in a 30-second window. For each trial, 0.2 mL Alcohol (20% EtOH) was delivered for 6 seconds into the magazine for CS+ trials.

Trials were followed by a variable inter-trial-interval (vITI 2mins). We trained the rats for 5 sessions in this phase.

##### Appetitive-Aversive Conflict Phase

Sessions were procedurally identical to the conditioning sessions, with the exception that a mild footshock (0.3mA intensity, 0.5s duration) was delivered at the end of CS+ presentation, randomly on 50% of CS+ trials. Shock delivery was determined using the random selection function (RANND) from the following array (0,1,0,1,0,1,0,1). We trained the rats for 4 sessions in this phase.

#### Exp. 2: Monitoring LH-GABA neurons during discrimination learning between appetitive and aversive conditioning

##### Home-cage alcohol phase

In a similar manner to previous work (Alonso-Lozares et al., 2024), we first habituated rats to alcohol consumption by using an alcohol homecage procedure (Wise, 1973; Simms et al., 2008). Rats had intermittent 24h access to a bottle containing 20% ethanol (EtOH), and a bottle containing water. Additionally, we used two control empty cages without animals to account for spillage. After 24h the bottles were weighed and swapped for a single water bottle for the following 24h. These two phases were repeated 12 times. The overall consumption of alcohol per rat after each session was measured in grams, by subtracting pre and post measurements for each cage and the average of the two control cages, accounting for 2-gram spillage and multiplying by the density of 20% ethanol (0.97).

##### Reward only phase

We trained the rats to associate a cue with alcohol delivery in the alcohol magazine (CSa trials). Cues consisted of two of three different stimuli (white noise, light, or flickering light) counterbalanced across rats. In each session, rats received 8 presentations of the CSa. Data were divided into three phases (pre-CS, during-CS, and post-CS), each lasting 20 seconds and resulting in a 60-second window. For each trial, 0.2 mL Alcohol (20% EtOH) was delivered over 6 seconds into the magazine during the trials starting from 14s after cue onset. Trials were followed by a variable inter-trial-interval (vITI 2mins). We trained the animals until each had stable behaviour (More than 3 consecutive sessions with > 3 seconds spent in the magazine during cue presentation), resulting in an unequal number of training sessions per animal (Range = 19 – 30 sessions). We only recorded the first two (Early) and last two (Late) session in this phase.

##### Appetitive-Aversive Discrimination Phase

We delivered one cue (CSs) that co-terminated with a footshock (0.3mA intensity, 0.5s duration), another cue with no programmed consequences (CS-), and the same CSa from the previous phase was also presented. In each session, rats received 8 presentations of each CS, randomly determined, with a total of 24 trials. Rats received 8-12 training sessions.

##### Appetitive-Aversive Conflict Phase

There were three trial types: Reward (CSa and CS-) which resulted in a 6 second alcohol delivery 14 seconds after the start of the cue period, Shock (CSs and CS-) which resulted in 0.5 sec shock (0.3 mA) when the cues are turned off, and Conflict (CSa and CSs) which resulted in 6 second alcohol delivery 14 seconds after the start of the cue period and a 0.5 mA shock (0.3 mA) when the cues are turned off. In each session, rats received 8 presentations of each compound cue type, randomly determined, with a total of 24 trials. Time spent in the magazine was defined in the same way as the previous phases. Rats were given 4-11 training sessions.

### Histological validation

#### Histological validation after behavioural experiments

After each behavioural experiment, we deeply anesthetized rats with isoflurane and Euthasol® injection (i.p.) and transcardially perfused them with ∼100 ml of normal saline followed by ∼400 ml of 4% paraformaldehyde (PFA) in 0.1M sodium phosphate (pH 7.4).

The brains were removed and post-fixed in PFA for 2 h, and then in 30% sucrose in 0.1M PBS for 48 h at 4°C. Brains were then frozen on dry ice, and coronal sections were cut (40 µm) using a Leica Microsystems cryostat and stored frozen in 0.1M PBS containing 30% sucrose at -20°C.

We selected a 1-in-4 series and first rinsed free-floating sections (3 x 10 minutes) before washing in PBS. Following this, slices were stained with DAPI (0.1 ug/ml) for 10 min prior to washing and mounting onto gelatin-coated glass slides, air-drying and cover-slipping with Mowiol and DABCO.

### Image acquisition, fiber placement, and jGCaMP7f expression verification

We used a VectraPolaris slide scanner (VUmc imaging core) to image slides at 10x magnification. Images from Bregma -2.04 mm to Bregma -2.76 mm were scanned and imported into QuPath (Bankhead et al., 2017). LH was identified using DAPI for identification of anatomical landmarks and boundaries, and the boundary of jGCaMP7f expression and fiber placement for each rat was plotted onto the respective atlas plate (Paxinos and Watson, 2008).

### Statistics

#### Behavioural data

We analysed magazine time (s) using GraphPad Prism 9.1.0. Phases were analysed separately. In experiment 1, the dependent variables were the average time spent in the magazine pre cue (10s), during cue (10s), and after cue (10s) for each cue type presented across the phases (Retraining, Shock). We used a repeated measures analysis of variance (ANOVA), with type of CS (CS+, CS-) and session (Retraining 1 vs. Retraining 5; Shock 1 vs. Shock 4) as the within-subject factors. In experiment 2, the dependent variables were the average time spent in the magazine pre cue (20s), during cue (20s), and after cue (20s) for each cue type presented in each phase. Specifically, Reward phase: CSa only; Discrimination phase: CSa, CS-, CSs; Conflict phase: Reward (CSa/CS-), Shock (CSs/CS-), Conflict (CSa/CSs). We used a repeated measures ANOVA, with type of CS as the within-subject factor, and follow-up posthoc tests to compare magazine entries between each cue type.

#### Photometry

Recorded signals were first down sampled by a factor of 64, giving a final sampling rate of 15.89 Hz. The 405 nm isosbestic signal was fit to the 490 nm calcium-dependent signal using a first-order polynomial regression. A normalized, motion-artifact-corrected ΔF/F was then calculated as follows: ΔF/F = (490 nm signal − fitted 405 nm signal)/fitted 405 nm signal. The resulting ΔF/F was then detrended via a 90-s moving average, and low-pass filtered at 3 Hz. ΔF/F from 10s before CS (baseline) to 20s after CS onset were collated.

These traces were then baseline corrected and converted into z-scores by subtracting the mean baseline activity during the first 5s of the baseline and dividing by the standard deviation of those 5s.

For experiment 1, traces were grouped by type of CS (CS+ or CS-) and by session (Retraining ‘Early’ (sessions 1 and 2) vs. Retraining ‘Late’ (sessions 4 and 5); Shock 1 vs. Shock 5). For experiment 2, traces were grouped by the experimental phase (Reward only, Discrimination, Conflict), and the cue type.

Both bootstrapping and permutation test analysis approaches were used, and the rationale for each is described in detail previously but will briefly be described below (Jean-Richard-Dit-Bressel et al., 2020; Ghareh et al., 2022). Bootstrapping was used to determine whether calcium activity during CS was significantly different from baseline (ΔF/F = 0). A distribution of bootstrapped means was obtained by randomly sampling from traces with replacement (n traces for that response type; 5000 iterations). A 95% confidence interval was obtained from the 2.5th and 97.5th percentiles of the bootstrap distribution, which was then expanded by a factor of sqrt (n/(n − 1)) to account for narrowness bias (Jean-Richard-Dit-Bressel et al., 2020).

Permutation tests were used to assess significant differences in calcium activity between CS types across the different phases of the tasks. Observed differences between CS type or session were compared against a distribution of 1000 random permutations (difference between randomly regrouped traces) to obtain a p-value per time point. An alpha of 0.05 was Bonferroni-corrected based on the number of comparison conditions, resulting in alpha of 0.01. For both bootstrap and permutation tests, only periods that were continuously significant for at least 0.25 s were identified as significant (Jean-Richard-Dit-Bressel et al., 2020).

The output of every statistical test we conducted is available in the raw data files. The significant outputs of these tests are indicated in the figures, below the traces. Specifically, the presence of the bar within the graph indicates the time periods in which the statistical test produced a significant output. For additional information, we have also reported the time periods in Tables 1 and 3, and therefore these values are not reported in the results section.

## Results

### Exp. 1: Activity of LH-GABA neurons during reacquisition of a cue-alcohol association, and during appetitive-aversive conflict conditioning

In experiment 1, we aimed to describe LH-GABA activity during reacquisition of a previously extinguished cue-alcohol association, and how LH-GABA activity integrates aversive information by introducing appetitive-aversive conflict conditioning to this cue. Figure 1A shows the experimental timeline, the expression of the Ca2+ sensor jGCaMP7f (Dana et al., 2019) in LH-GABA was achieved using a mixture of one AAV encoding Cre driven by the GAD promoter with another AAV encoding Cre-dependent jGCaMP7f under the hSyn1 promoter. We have previously validated this viral targeting approach (Alonso-Lozares et al., 2024). Figure S1 shows the experimental timeline prior to experiment 1.1 in this study, and the corresponding behavioural data reported in Alonso-Lozares et al., (2024). We measured fluorescence via an optic fiber cannula implanted above LH throughout the entire experiment (Figure 1B).

**Figure 1.**
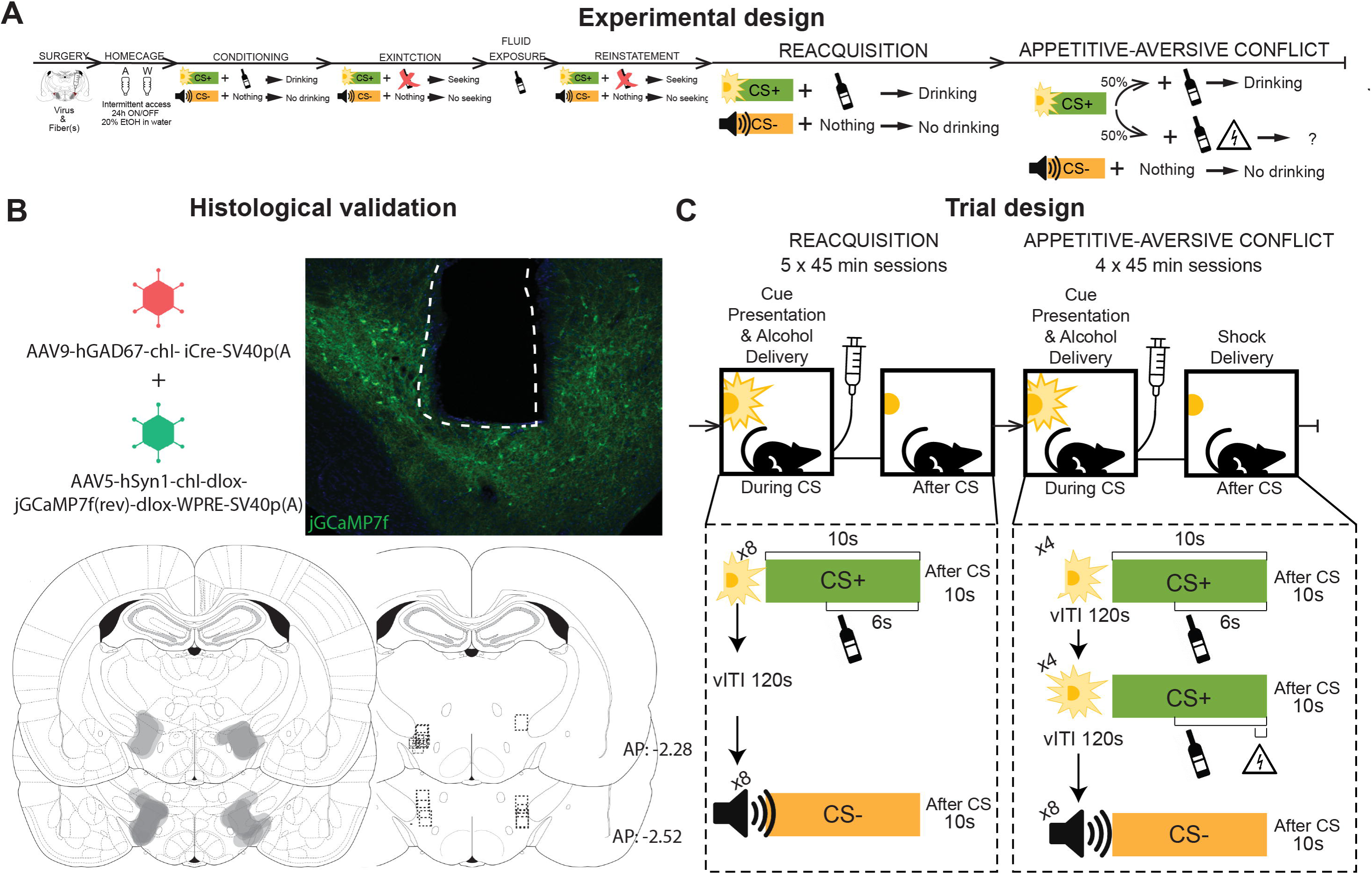
Monitoring LH-GABA neurons during reacquisition of a previously learned cue-alcohol association, and appetitive-aversive conflict. (**A**) Outline of the experimental procedure (n = 13 females). See Figure S1 for the procedure between home-cage alcohol and reacquisition. (**B**) Representative image of jGCaMP7f expression and fiber implant in LH, and fiber placement and expression validation. (**C**) Trial design for the experiment.

### Experiment 1.1: Reacquisition of a previously extinguished cue-alcohol association

#### Behavioral data

We used repeated measures ANOVA with the within-subjects factors Session (Early, Late) and CS Type (CS+, CS-) to compare magazine entries during the reacquisition sessions (Figure 2A and 2B). We found main effects of Session and CS Type, as well as a significant Session x CS Type interaction on time spent in the magazine during the cue period (Figure 2B; F (1, 12) = 26.8, p < 0.001), but not after the cue period (Figure 2C; F (1, 12) = 4.697, p = 0.051). We found no difference between CS Type on time in the magazine during the pre-CS time period (Figure S2). These data show that, throughout the course of the reacquisition phase, rats successfully discriminate between the alcohol-predictive and neutral cues, and that time in the magazine during the cue period increases across the reacquisition sessions.

**Figure 2.**
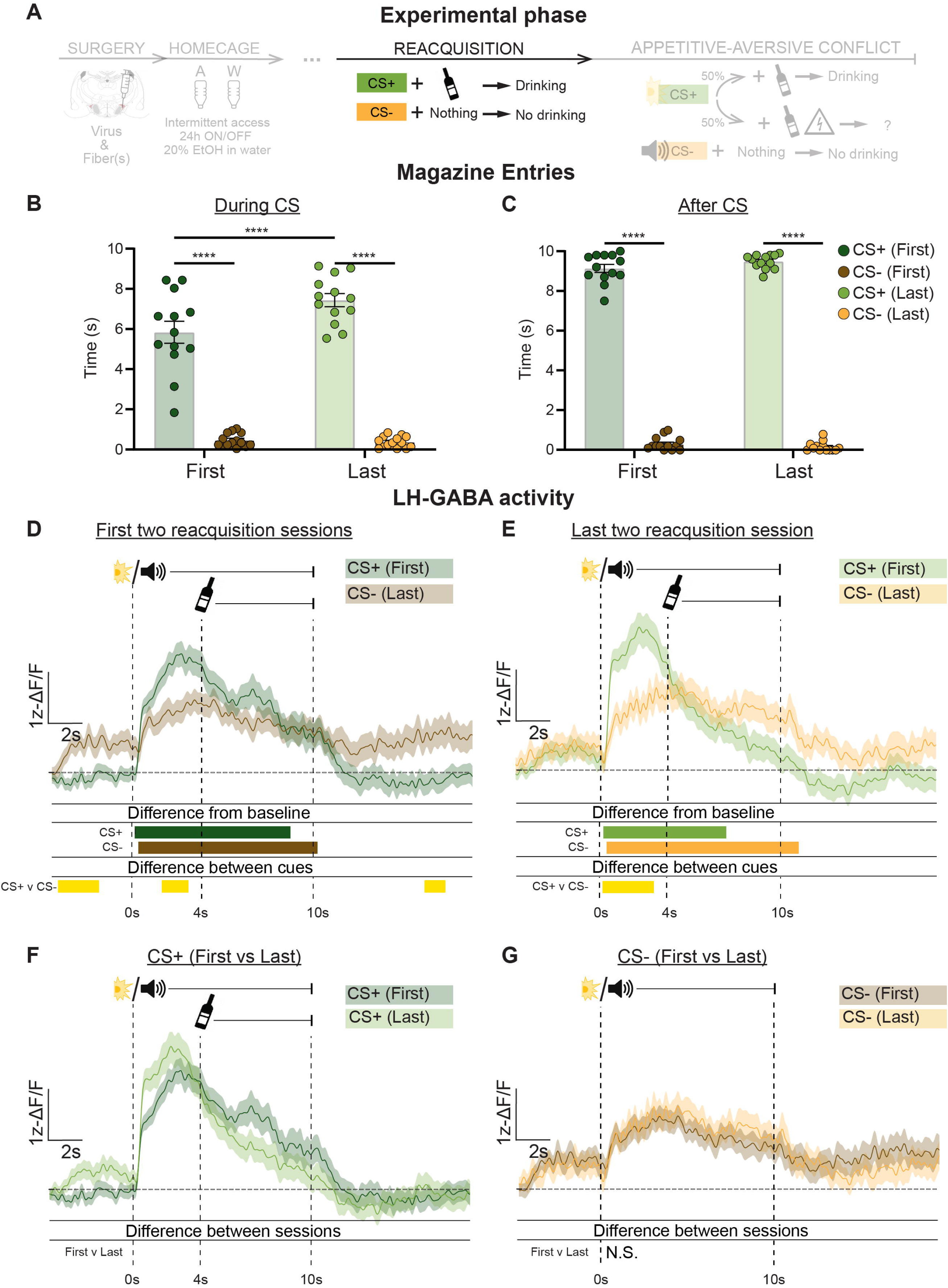
Monitoring LH-GABA neurons during reacquisition of a previously extinguished cue-alcohol association. (**A**) Outline of experimental procedure (n = 13 females). Mean ± standard error of the mean (SEM) time spent in the alcohol magazine during **(B)** and after **(C)** the cue period comparing the first and last reacquisition sessions (n = 8 trials per CS type, per session). Alcohol was delivered at 4s after CS onset. Ca2+ traces of LH-GABA activity centred around cue onset (−5s to +20s) comparing LH-GABA activity to CS+ and CS-on the early **(D)** and late **(E)** reacquisition sessions. The same mean Ca2+ traces are plotted for comparisons across sessions between CS+ **(F)** and CS-**(G)**. For all photometry traces, bars at bottom of graph indicate significant deviations from baseline (dF/F ≠ 0) determined via bootstrapped confidence intervals (99% CIs), or significant differences between the specific events (CS+ early; CS-early; CS+ late; CS-late) determined via permutation tests with alpha = 0.01 for comparisons between CS type and session. Vertical dashed lines indicate CS onset and alcohol delivery for CS+ trials, horizontal line indicates baseline (dF/F = 0).

#### Photometry data

We made within-session comparisons (Figures 2D, 2E) between the CS types (CS+ vs CS-), and across-session comparisons (Figures 2F, 2G) within the same CS type (Early vs. Late). Detailed significance time windows can be found in Table 1. Figure 2D shows LH-GABA activity recorded in the early reacquisition sessions. We found that LH-GABA activity significantly increased from baseline to both CS types but was significantly higher during the CS+ compared to the CS- (permutation tests) both during cue onset and after alcohol delivery. Figure 2E shows LH-GABA activity recorded in the late reacquisition sessions. Here, activity is significantly increased to both CS types, and there were significant differences between the cues after cue onset, and after alcohol delivery. These data show that reacquisition of a previously extinguished cue-alcohol memory is associated with increased activity of LH-GABA neurons to the alcohol-predictive cue, and that the pattern of activity to CS+ and CS- is comparable in this phase to before extinction (Alonso-Lozares et al., 2024).

To further investigate whether LH-GABA activity changes across the sessions, we performed additional analyses making across-session comparisons within the same CS type. Here, permutation tests revealed no significant differences in LH-GABA activity for either CS+ (Figure 2F) or CS- (Figure 2G) across the sessions. These analyses show that reacquisition of the cue-alcohol association was associated with increased activity in LH-GABA neurons to the alcohol predictive CS+ from the first session, and that this pattern of activity remains constant across reacquisition sessions.

### Experiment 1.2: Appetitive-aversive conflict sessions

#### Behavioral data

We used repeated-measures ANOVA with the within-subjects factors Session (First, Last) and CS Type (CS+, CS-) to compare magazine entries during this phase (Figure 3A). Figure 3B shows time spent in the magazine during the cue period, and Figure 3C shows time spent in the magazine after the cue period. We found main effects of Session and CS Type, as well as a significant Session x CS Type interaction on time spent in the magazine during (F (1, 12) = 101.0, p < 0.0001), and after (Figure 3C; F (1, 12) = 15.85, p = 0.01) the cue period. We found no difference between CS Type on time in the magazine during the pre-CS time period (Figure S3). These data show that introduction of a potential shock outcome significantly reduces magazine entries during the cue period, however magazine entries after the cue period remains significantly higher for the CS+ than the CS-, despite the foot shock.

**Figure 3.**
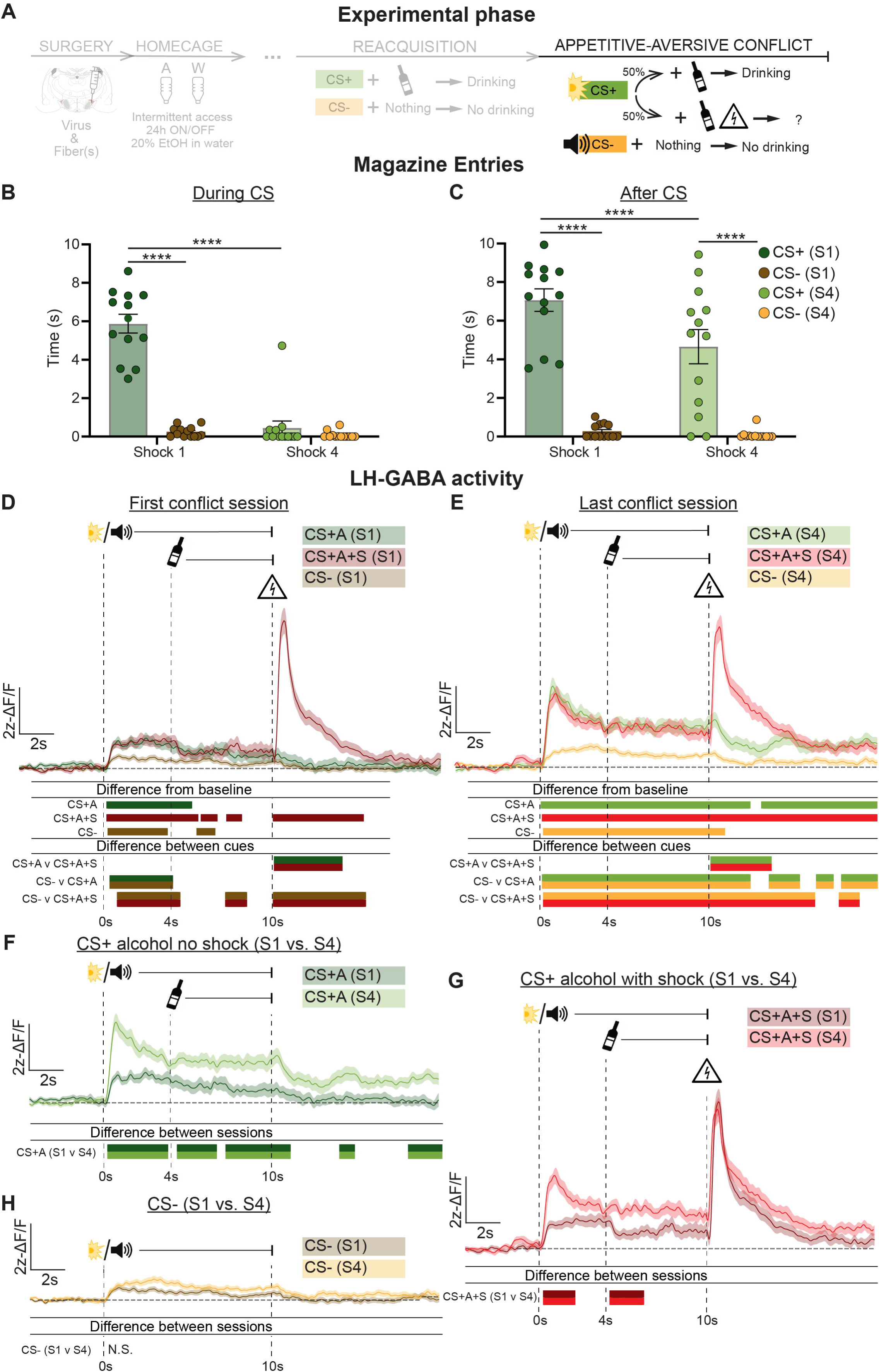
Monitoring LH-GABA neurons during appetitive-aversive conflict. (**A**) Outline of experimental procedure (n = 13 females). Mean ± standard error of the mean (SEM) time spent in the alcohol magazine during **(B)** and after **(C)** the cue period comparing the first and last shock sessions (n = 8 trials per CS type, per session). Alcohol was delivered at 4s after CS+ onset, footshock was delivered after CS+ offset for 0.5s on 50% of CS+ trials. Ca2+ traces of LH-GABA activity centred around cue onset (−5s to +20s) comparing LH-GABA activity to CS+ and CS- on the first **(D)** and last **(E)** shock sessions. The same mean Ca2+ traces are plotted for comparisons across sessions between CS+ with alcohol only **(F)** CS+ with alcohol and shock **(G)**, and CS- **(H)**. For all photometry traces, bars at bottom of graph indicate significant deviations from baseline (dF/F ≠ 0) determined via bootstrapped confidence intervals (99% CIs), or significant differences between the specific events (CS+ Shock1; CS-Shock1; CS+ Shock4; CS-Shock4) determined via permutation tests with alpha = 0.01 for comparisons between CS type and session. Vertical dashed lines indicate CS onset and alcohol delivery for both CS+A and CS+A+S trials, and shock delivery for CS+A+S trials. Horizontal line indicates baseline (dF/F = 0).

#### Photometry data

We made within-session comparisons (Figures 3D, 3E) between the CS types (CS+A, CS+A+S, CS-), and across-session comparisons (Figures 3F, 3G, 3H) within the same CS type (First vs. Last). Figure 3D shows LH-GABA activity from the first appetitive-aversive conflict session. Detailed significant time windows can be found in Table 1. We found that LH-GABA activity was significantly higher than baseline for all CS types: CS+A (alcohol only), CS+A+S (alcohol+shock), and CS-. Permutation tests revealed differences between CS+ and CS-during the initial cue period regardless of the final outcome, and that LH-GABA activity is substantially increased after the shock.

Figure 3E shows LH-GABA activity recorded in the last appetitive-aversive conflict session. We also find here, using bootstrapped CI tests, that LH-GABA activity significantly increased from baseline to all CS types. Permutation tests revealed differences between both CS+ conditions and CS-during the initial cue period, and we also found significant differences in LH-GABA activity to the shock.

To show how LH-GABA neuronal activity changes with appetitive-aversive conflict learning, we performed additional analyses comparing LH-GABA activity between the first and last sessions (Figure 3 F-H). Permutation tests revealed significant differences in the initial cue period between the first and last sessions for both CS+ conditions, reflecting higher activity to the CS+ in session 4 compared to session 1. We found no difference between first and last sessions in LH-GABA activity associated with the CS-. Finally, there was no significant difference between session 1 and session 4 after the shock. Overall, these data show that when a cue which predicts alcohol changes to also predict potential aversive outcomes, LH-GABA activity significantly increases to this cue. Furthermore, we found no change in LH-GABA activity to the shock outcome across learning (Figure 3G), indicating that the salience of the shock outcome itself does not change in this experimental procedure.

### Experiment 2: Activity of LH-GABA neurons during discrimination learning between appetitive and aversive conditioning

In experiment 2, we aimed to describe LH-GABA activity during acquisition of a novel cue-alcohol association, followed by acquisition of a novel cue-shock association. We also tested how LH-GABA activity integrates appetitive and aversive information in conflict tests where these cues are presented in compound. Figure 4A and 5A show the experimental timeline, and we measured fluorescence via an optic fiber cannula implanted above LH throughout the entire experiment (Figure 4B).

**Figure 4.**
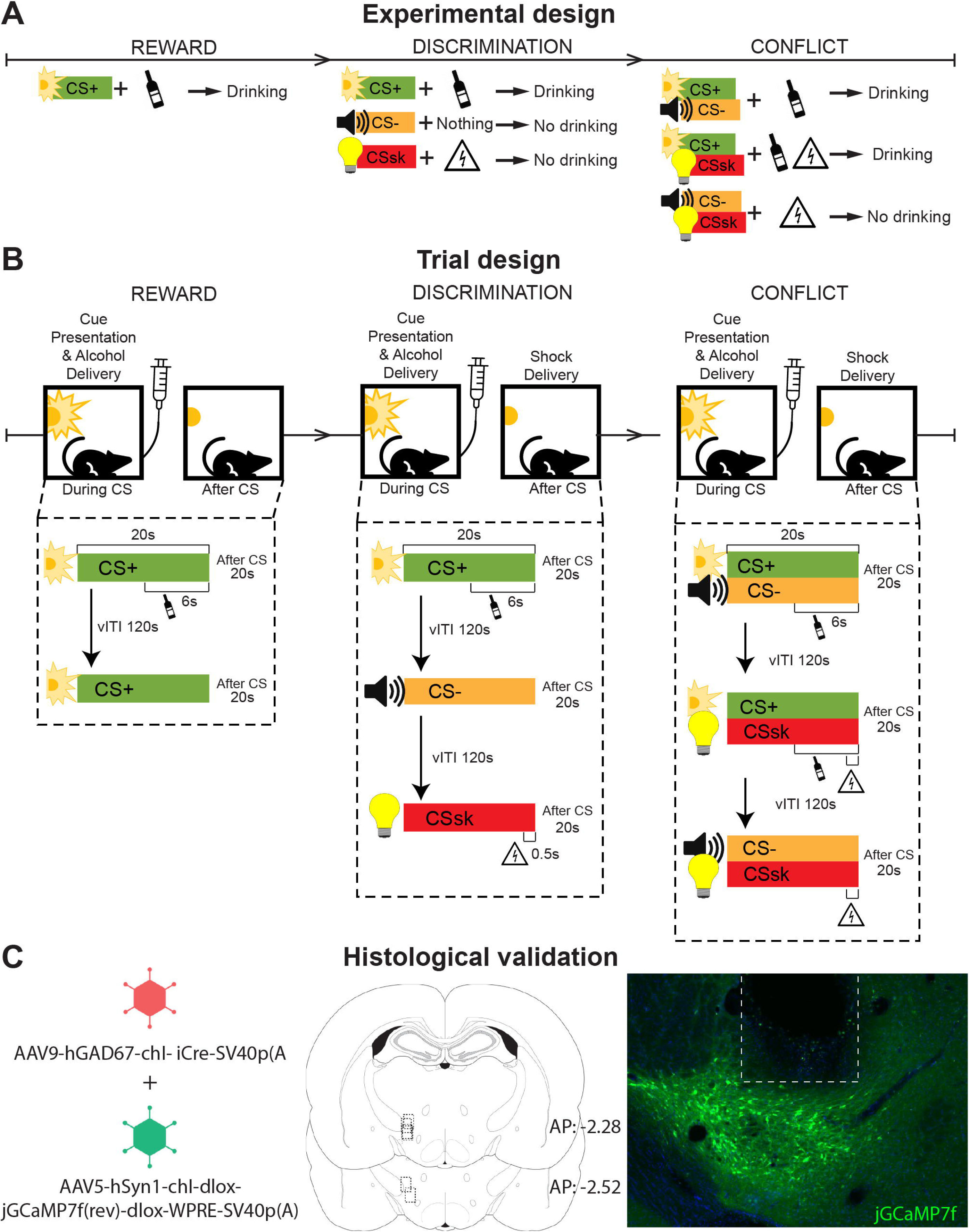
Pavlovian conflict task design. (**A**) Outline of the experimental procedure (n = 13 females). (**B**) Trial design for each phase of the experiment (n = 8 trials per CS type, per session, randomly presented). Alcohol was delivered at 14 s after CS+ onset for Reward, and Appetitive-Aversive Discrimination sessions, and 14s after Reward and Conflict trials CS onset for Conflict sessions. Foot shock was delivered after CSs offset for 0.5s in Discrimination sessions and Conflict sessions. (**C**) Representative image of jGCaMP7f expression and fiber implant in LH, and fiber placement and expression validation for rats in experiment 2.

**Figure 5.**
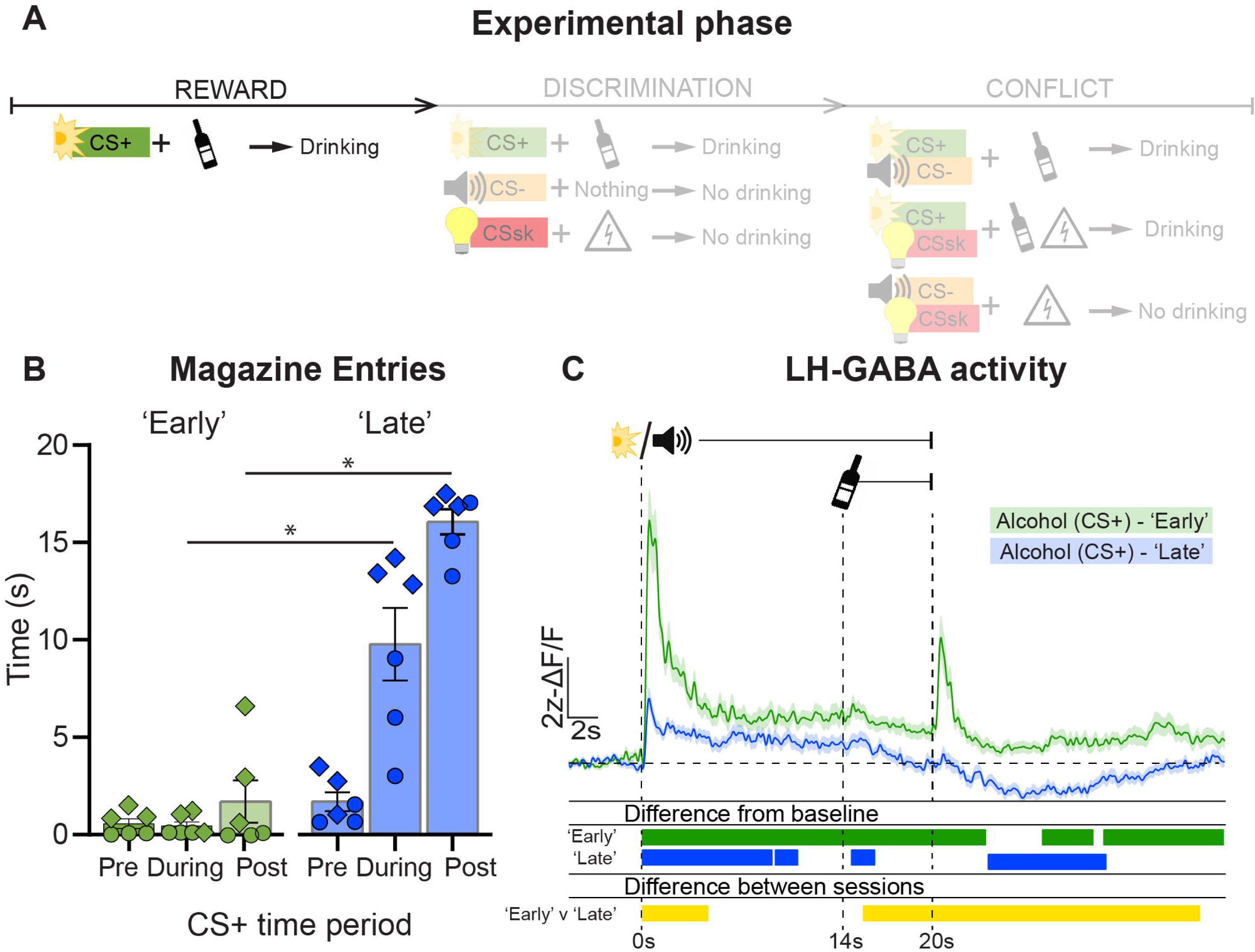
Monitoring LH-GABA neurons during reward training. (**A**) Outline of experimental procedure (n = 3 males; diamond shapes, n = 3 females, circle shapes). **(B)** Mean ± standard error of the mean (SEM) time spent in the alcohol magazine before, during and after CSa presentation across all reward sessions (n = 8 trials per session). **(C)** Ca2+ traces of LH-GABA activity centred around cue onset (−5s to +40s) showing activity to CSa onset across all reward sessions. Alcohol was delivered at 14 s after CS onset. Bars at bottom of graph indicate significant deviations from baseline (dF/F ≠ 0) determined via bootstrapped confidence intervals (99% CIs). Vertical dashed lines indicate CS onset and alcohol delivery. Horizontal line indicates baseline (dF/F = 0).

### 2.1 Monitoring LH-GABA neurons during cue-alcohol association acquisition

#### Behavioral data

We used repeated-measures one-way ANOVAs with the within-subjects factors Session (Early, Late) and Time Period (Pre, During, Post) to compare magazine entries between before, during, and after CSa onset from the first two to last two conditioning sessions (Figure 5B). We found a Session x cue period interaction (F (2, 10) = 31.4, p < 0.001). Further analyses (Table 2) show significant differences in magazine entries between the ‘during’ and ‘post’ cue time periods, comparing early and late.

#### Photometry data

Figure 5C shows LH-GABA activity during cue presentations in the Reward sessions. Detailed significant time windows can be found in Table 3. We found significantly increased LH-GABA activity relative to baseline to the alcohol predictive cue, during alcohol delivery and at the offset of the alcohol predictive cue. These data show, like we observe in Exp. 1, that LH-GABA activity is increased to an alcohol-predictive cue.

### Experiment 2.2 Appetitive-Aversive Discrimination training

#### Behavior

We used repeated-measures one-way ANOVAs with the within-subjects factor CS Type (CSa, CS-, CSs; Figure 6B) to compare magazine entries during and after CS onset (Figure 6B and 6C). We found a main effect of CS Type during the cue period F (1.8, 8.8) = 6.8, p < 0.05), but no significant post-hoc differences between the individual CS types, although the trend shows rats spent more time in the magazine during CS+ presentation (Table 2). We also found a main effect on magazine entries after the cue period (F (1.3, 6.6) = 81.9, p < 0.001), as well as significant interaction between CSa and CS-, and CSa and CSs (Table 2). We found no difference between CS Type on time in the magazine during the pre-CS time period (Figure S4). These data show that the rats discriminate between the different cue types, and that magazine entries are increased during CS+ trials, but not the other types.

**Figure 6.**
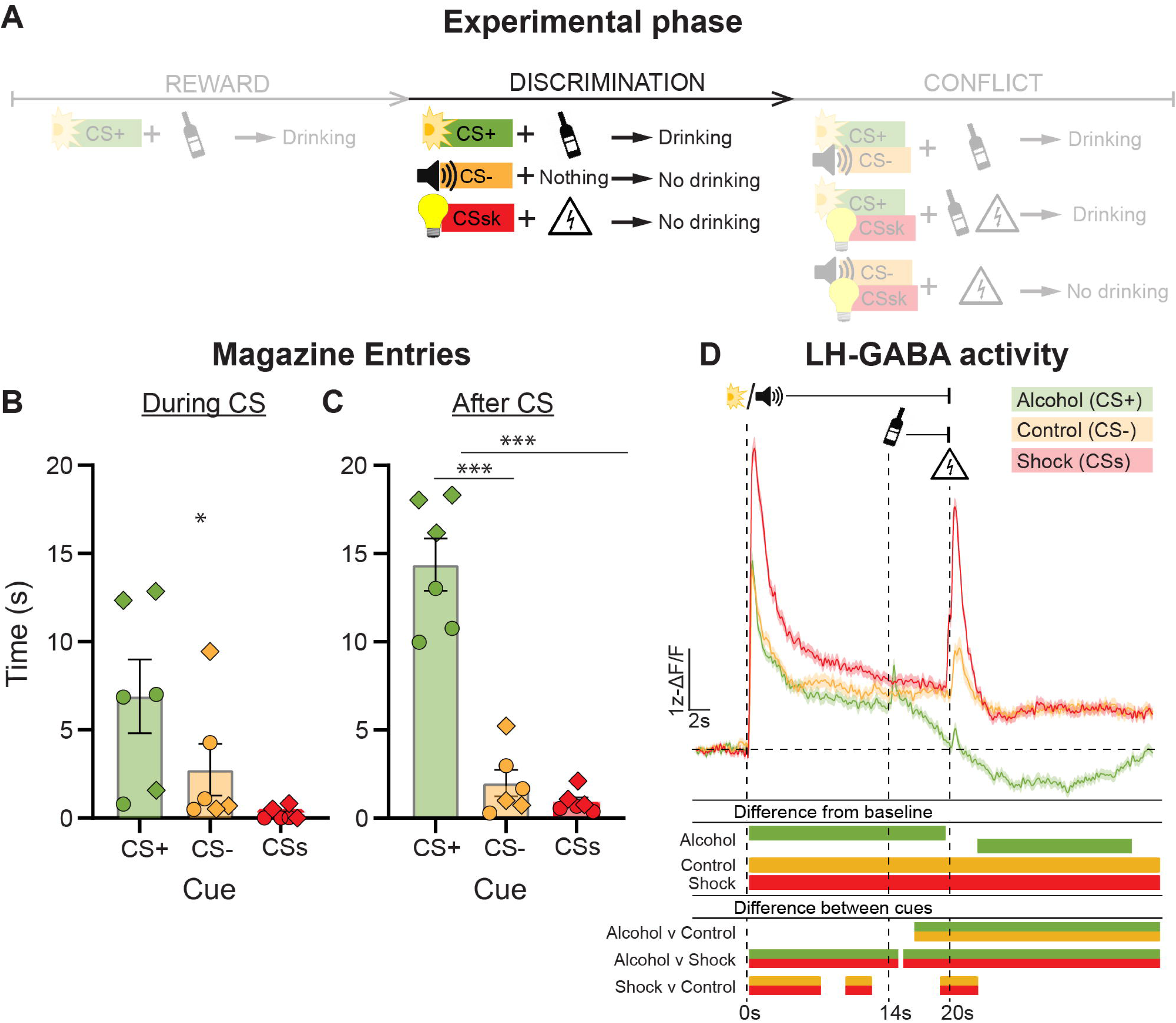
Monitoring LH-GABA neurons during discrimination training. (**A**) Outline of experimental procedure (n = 3 males; diamond shapes, n = 3 females, circle shapes). Mean±SEM time spent in the alcohol magazine during **(B)** and after **(C)** the cue period during discrimination sessions (n = 8 trials per CS type, per session). Alcohol was delivered at 14 s after CS+ onset, foot shock was delivered after CSs offset for 0.5s. **(D)** Ca2+ traces of LH-GABA activity centred around cue onset (−5s to +40s) comparing LH-GABA activity to CS+, CS-, and CSs on all discrimination sessions. Bars at bottom of graph indicate significant deviations from baseline (dF/F ≠ 0) determined via bootstrapped confidence intervals (99% CIs), or significant differences between the specific events (CSa; CS-; CSs) determined via permutation tests with alpha = 0.01 for comparisons between CS type. Vertical dashed lines indicate CS onset, alcohol delivery for CSa trials, and shock delivery for CSs trials. Horizontal line indicates baseline (dF/F = 0).

#### Photometry

We made comparisons between the CS types (CSa vs CS- vs CSs) across all sessions in the Discrimination phase (Figure 6D). Detailed significant time windows can be found in Table 3. We found that LH-GABA activity was significantly higher for all three cue types during the cue period in the Discrimination phase. Bootstrapping analysis revealed a significant increase from baseline for CSa followed by a significant decrease from baseline after alcohol delivery. We also observed a significant increase from baseline and throughout the entirety of the trial for both CS- and CSs. Permutation tests revealed differences in LH-GABA activity in the cue period between CSs and CSa, and between CSs and CS-, reflecting significantly higher LH-GABA activity to the shock-predictive cue than the other two cue types. During the post-cue period, we found that LH-GABA activity is significantly higher to the shock outcome compared to alcohol after the CSa, and to nothing after the CS-.

These data show that LH-GABA neurons activity is higher to a shock predictive cue compared to an already-established alcohol-predictive cue. Both CS- and CSs were introduced in the same experimental phase, which may be why CS-induced activity is comparable to the CSa, but significantly lower than to the shock predictive cue.

### Experiment 2.3: Conflict training

In this phase, we presented the three previously conditioned cues in three compound cue pairs. There were three compound cue types: Reward (CSa/CS-), Shock (CSs/CS-), and Conflict (CSa/CSs), and the following outcomes: Reward → alcohol; Shock → Footshock; Conflict → Alcohol+Footshock.

#### Behavior

We used repeated-measures one-way ANOVAs with the within-subjects factors compound CS Type (Reward, Shock, Conflict) to compare magazine entries during the cue period (Figure 7B) and after the cue period (Figure 7C). We found a main effect of Cue Type during the cue period (F (1.0, 5.1) = 8.3, p < 0.05), but no post-hoc differences. For magazine entries after the cue period, we found a main effect of Cue Type (F (1.5, 7.4) = 13.6, p < 0.01). Post-hoc analysis revealed significant differences between Conflict and Reward Trials, and Conflict and Shock trials (Table 2). We found no effect of Cue Type on time in the magazine during the pre-CS time period (Figure S5). These data show that the trial types where the shock-predictive cue is presented (Shock or Conflict) leads to significantly less magazine entries during the cue period compared to when the shock-predictive cue is not presented (Reward). However, after the cue period magazine entries are not different between the Reward and Conflict trials, and both are significantly higher than the Shock trials when no alcohol was delivered.

**Figure 7.**
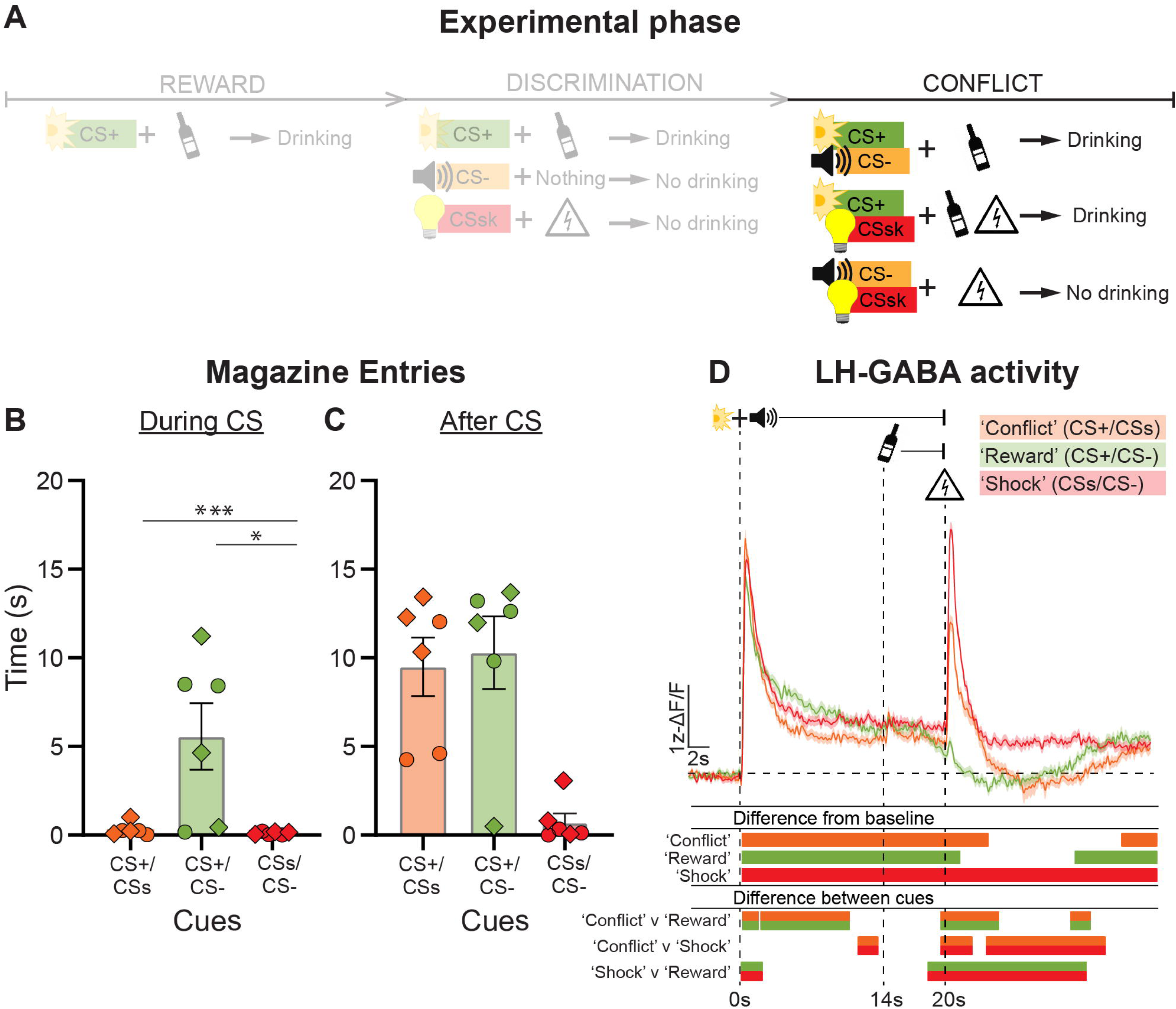
Monitoring LH-GABA neurons during the conflict phase. (**A**) Outline of experimental procedure (n = 3 males; diamond shapes, n = 3 females, circle shapes). Mean ± standard error of the mean (SEM) time spent in the alcohol magazine during **(B)** and after **(C)** the cue period during Conflict sessions (n = 8 trials per CS type, per session). Alcohol was delivered at 14 s after Reward and Conflict trial CS onset, foot shock was delivered after Shock and Conflict trial CS offset for 0.5s. **(D)** Ca2+ traces of LH-GABA activity centred around cue onset (−5s to +40s) comparing LH-GABA activity to Conflict, Reward and Shock on all Conflict sessions. Bars at bottom of graph indicate significant deviations from baseline (dF/F ≠ 0) determined via bootstrapped confidence intervals (99% CIs), or significant differences between the compound CS determined via permutation tests with alpha = 0.01 for comparisons between compound CS type. Vertical dashed lines indicate CS onset, alcohol delivery for Reward and Conflict trials, and shock delivery for Shock and Conflict trials. Horizontal line indicates baseline (dF/F = 0).

#### Photometry

We made comparisons between the compound cue types (Conflict vs Reward vs Shock) across all recorded sessions in the Conflict phase (Figure 7D). Detailed significant time windows can be found in Table 3. We found, using bootstrapped CI, that LH-GABA activity was significantly increased from baseline for all three compound cue types during the cue period in the conflict phase. Permutation tests revealed significant differences in LH-GABA activity during the cue period between Conflict and Reward trials and between Shock and Reward trials. This shows that cue-induced LH-GABA activity is higher in the presence of the shock-predictive cue (CSs), as both trial types that contained this cue were higher than the other trial type (Reward) which did not contain this cue. Finally, we also found significant differences in LH-GABA activity in response to the shock between the trial types. Permutation tests revealed significantly lower LH-GABA activity to the shock during Conflict trials compared to Shock trials, and significantly higher LH-GABA activity during the Conflict trials compared to the Reward trials. These data suggest an interaction of appetitive (alcohol) and aversive (shock) encoding in the conflict trials, such that alcohol reduces the degree of activation caused by the shock in this experimental procedure.

## Discussion

Here we describe how LH-GABA neurons encode reward and fear learning, and how this activity changes during conflict. We first show that activity to a reward-predictive cue increases when that cue predicts footshock. We next trained a different group of rats to associate distinct cues with different appetitive or aversive outcomes and found that activity is highest for the shock-predictive cue. When these cues are presented in compound, cue-induced activity was highest for the compound cues containing the shock cue. During outcome delivery however, we found that shock-induced activity was attenuated when shock was delivered with alcohol compared to without alcohol. These data show that activity tracks both conditioned and unconditioned salient events. We found no evidence for attenuated activity to a shock-predictive cue during conflict, however activity was attenuated to the shock outcome during conflict. Attenuation of aversive encoding in the presence of alcohol may be a potential mechanism of continued alcohol use despite negative consequences.

### Methodological considerations

Several issues must be considered in the interpretation of these findings. Firstly, rats in experiment 1 had a history of extinction prior to the appetitive-aversive conditioning. The impact of this prior learning on our findings is limited because the reacquisition data shows a pattern, both in behaviour and LH-GABA activity, that is comparable to prior to extinction (Alonso-Lozares et al., 2024). The effect of sex and oestrous cycle on the neurobiological underpinnings of behaviour has gained interest in recent years (Torres et al., 2014). In experiment 1, we only used female rats and did not measure oestrus cycle. While we cannot exclude cycle effects in this experiment, recent work has shown no sex differences in operant behaviour related to drinking (Priddy et al., 2017). In experiment 2, we used both male and female rats and observed no significant differences in behaviour or LH-GABA activity. Therefore, we assume that LH-GABA neuron activity of male and female rats is comparable.

Finally, although our findings show a specific role of the GAD-expressing subpopulation of LH neurons in cue-alcohol reward learning, these are a heterogeneous population of neurons. LH-GABA neurons are comprised of populations which express different peptides (Mickelsen et al., 2017; Mickelsen et al., 2019), and have different output projections (Bonnavion et al., 2016). Therefore, in this study, we have recorded the LH-GABA sub-population comprised of various phenotypes and efferent targets. It will be of interest in future studies to determine which of these outputs are critical for alcohol reward and aversive learning, and how these may interact in approach-avoidance conflict.

### Evidence for LH-GABA activity related to attention and salience, rather than valence

Here we described how LH-GABA activity encodes competition between cues predicting appetitive and aversive outcomes. We found that cue-induced activity is highest for aversive-predictive cues. This is consistent with observations that LH activity is necessary for fear learning after acquisition of a reward association (Sharpe et al., 2021). While greater LH-GABA activity was observed for shock-predictive cues, both appetitive and aversive predictive cues increased activity relative to baseline. Therefore LH-GABA activity is more closely related to salience than valence, a function previously demonstrated for LH (Wheeler et al., 2014). An alternative possibility is that distinct populations of neurons encode appetitive or aversive outcomes, or the cues which predict them.

Previously, we used a two-cue design and found that across learning the main change in LH-GABA activity was decreased to the control cue, with no change to the alcohol-predictive cue (Alonso-Lozares et al., 2024). In this study, however, we show that LH-GABA activity decreases across learning. We propose that the presence of two cues requires greater attentional demand. Specifically, in the single cue design no further processing is required to implement the behavioral response. Combined, these findings indicate that LH-GABA activity dynamically encodes the motivational or attentional processing required by the learning context, rather than simply signaling reward prediction itself.

### Evidence for integration of appetitive and aversive motivational states in LH-GABA activity

In experiment 1, we observe comparable responses to the shock across five sessions, indicating no habituation of LH-GABA activity to the shock. However, in experiment 2 we show that LH-GABA neuron activity integrates rewarding and aversive outcomes. Experimental psychologists have proposed the existence of two mutually inhibitory systems responsible for approaching rewarding outcomes and avoiding aversive outcomes (Konorski, 1967; Dickinson and Dearing, 1979). There is evidence for distinct populations encoding valence in brain regions such as the amygdala (Namburi et al., 2015; Beyeler et al., 2016; Burgos-Robles et al., 2017), but to date there is no evidence for this in the function of LH-GABA neurons. The presence of both appetitive and aversive stimuli are proposed to have mutually inhibitory effects on learning (Dickinson and Pearce, 1977). This study indicates that LH-GABA neurons may be a neurobiological locus of this mechanism. Interestingly, in experiment 2, magazine entries during the cue in the conflict condition was substantially decreased relative to the reward condition, but there was no difference in magazine entries in the post-cue period. This shows that shock expectancy is sufficient to abolish alcohol seeking during the cue period, but that once the shock is experienced alcohol taking is unaffected.

Our finding that shock-induced LH-GABA activity was attenuated when shock was delivered with alcohol provides a neural substrate for the summation principle of the Rescorla-Wagner model (Rescorla and Wagner, 1972; Wagner and Rescorla, 1972). According to this model, when CSa and CSs are presented together, their associative strengths should algebraically summate (Yau and McNally, 2023). Our data show that the activity of LH-GABA to the cue presentation shows no evidence for summation, however we show that LH-GABA neurons may integrate the motivational significance of competing outcomes, in a manner that bears resemblance to the principle of summation. This is potential candidate for alcohol consumption despite negative consequences, where the aversive impact of negative outcomes is reduced in the presence of alcohol. This process could contribute to the persistence of alcohol use despite mounting negative consequences observed in alcohol use disorder (McDonald et al., 2024).

### Broader LH-GABA circuitry related to motivation and salience

Previous research suggests that both appetitive and aversive learning involves excitatory projections from basolateral amygdala (BLA) to LH (Reppucci and Petrovich, 2016). The BLA is involved in both appetitive (Blundell et al., 2003; Holland and Gallagher, 2003) and aversive (Parkes and Westbrook, 2010; Sengupta et al., 2018) learning by signaling salient sensory-specific properties of stimuli and their relationship with the outcome (Hoang and Sharpe, 2021). This information might then be relayed through reciprocal LH-GABA projections to VTA (Watabe-Uchida et al., 2012; Nieh et al., 2015; Nieh et al., 2016). VTA also projects to BLA and these dopaminergic projections contribute to fear learning (de Oliveira et al., 2011). The LH-GABA to VTA pathway is potentially critical for the acquisition of the cue-alcohol memory. In previous work, we showed that LH-GABA projections to VTA regulate learning about food reward-paired cues (Sharpe et al., 2017), thereby identifying this pathway as critical for learning. The BLA-LH-VTA circuit may be critical for responding to the initial sensory-specific properties of stimuli, relayed to the LH through its glutamatergic projections from BLA. Previous work has shown that LH is involved in arousal and attentional states (Gutierrez et al., 2011; Wheeler et al., 2014). Attention is also a critical component of learning (Pearce and Hall, 1980; Le Pelley et al., 2016), and psychiatric disorders marked by maladaptive responses to motivational conflict, such as addiction, are characterised by attentional biases to drug-related cues (Field and Cox, 2008; Anderson, 2016). Increased LH-GABA activity may contribute to learning through signaling a heightened attentional state.

### Potential relationship to psychopathology

Our findings have implications for understanding psychological disorders characterized by maladaptive conflict resolution. In substance use disorders, attenuation of shock-induced LH-GABA activity in the presence of alcohol may reflect a neurobiological mechanism for continued substance use despite negative consequences (Hopf and Lesscher, 2014). Disruption in value integration could contribute to the attentional bias toward drug cues (Field and Cox, 2008) and impaired response inhibition (Goldstein and Volkow, 2011; Zilverstand et al., 2018) that characterize addiction. Similarly, anxiety disorders involve heightened sensitivity to threat cues and conflict detection (Gray and McNaughton, 2000), which may relate to enhanced LH-GABA response. Mechanistically, LH-GABA neurons are positioned to influence conflict processing through connections described above, potentially modulating cognitive processes fundamental to adaptive conflict resolution such as attentional salience attribution, response inhibition, and value integration.

### Concluding remarks

Our results show that LH-GABA activity significantly changes in a learning-dependent manner for both alcohol reward, and aversive shock, associations. When presented in conflict, we found that the shock-predictive cue caused a greater increase in activity compared to an alcohol-predictive cue. Future studies are therefore needed to determine the extent to which this activity is functionally relevant to choose appropriate behaviours, and to shed light on the circuitry that might orchestrate this type of learning. Overall, our findings highlight an important role for LH-GABA neurons in a fundamental learning process and show that this system is a critical component of both aversive and appetitive memory formation.

## Acknowledgments

The authors gratefully acknowledge the VUmc Histology Imaging Unit for their support & assistance in whole-slide imaging. Dr. Philip Jean-Richard-Dit-Bressel for assistance with photometry analysis and sharing early versions of the scripts we used.

