## Supplemental Figures for "Dual Role of LH-GABA Neurons in Encoding Alcohol Reward and Aversive Memories"

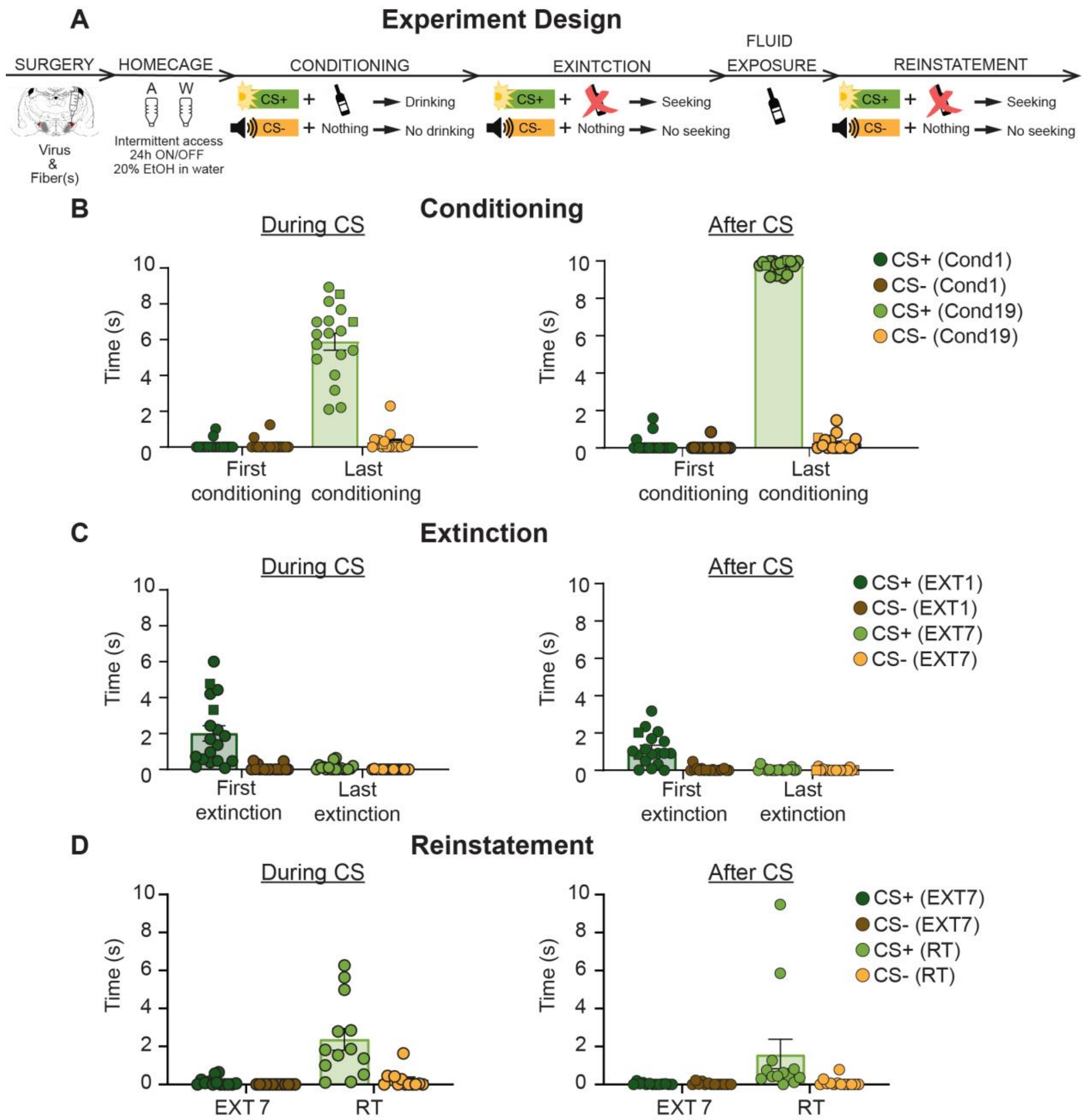


### Supplemental Figure Legends

**Figure S1. Experimental timeline of conditioning, extinction, and reinstatement prior to the first phase of Exp. 1.1.** (**A**) Experimental timeline. (**B, C, D**) Mean ± standard error of the mean (SEM) time spent in the alcohol magazine during CS and after the CS during the Conditioning (**B**), Extinction (**C**), and Reinstatement (**D**) phases of experiment 1, prior to Experiment 1.1 of this study. More details reported in Alonso-Lozares et al., (2024).

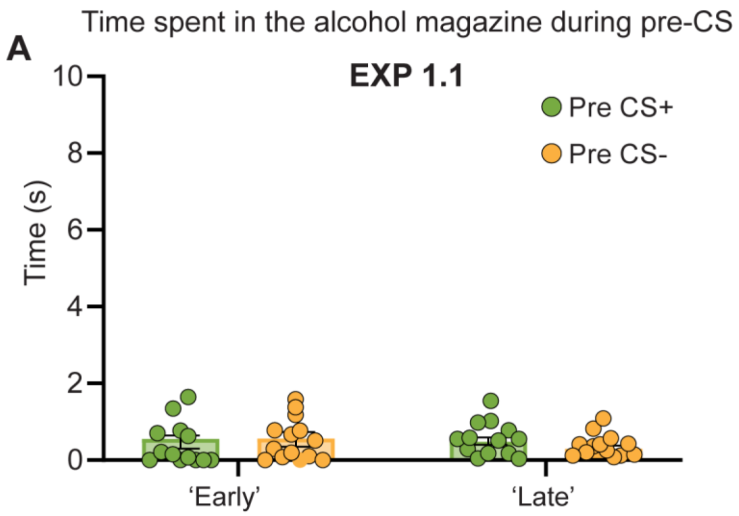


**Figure S2. Time spent in the alcohol magazine during pre-CS period in Exp. 1.1.** Mean ± standard error of the mean (SEM) time spent in the alcohol magazine during pre-CS in the reward phase of experiment 1.

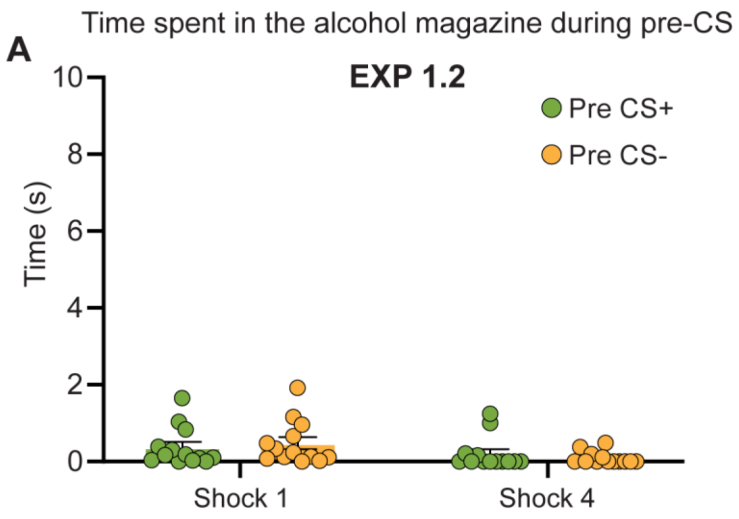


**Figure S3. Time spent in the alcohol magazine during pre-CS period in Exp. 1.2.** Mean ± standard error of the mean (SEM) time spent in the alcohol magazine during pre-CS in the conflict phase of experiment 1.

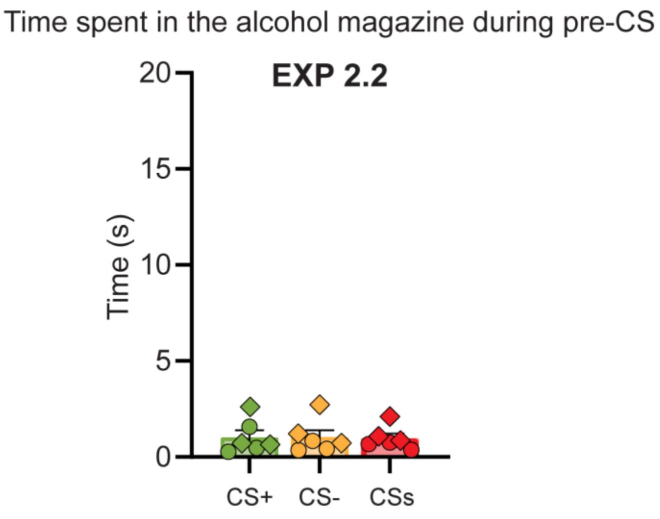


**Figure S4. Time spent in the alcohol magazine during pre-CS period in Exp. 2.2.** Mean ± standard error of the mean (SEM) time spent in the alcohol magazine during pre-CS in the discrimination phase of experiment 2. (n = 3 males; diamond shapes, n = 3 females, circle shapes).

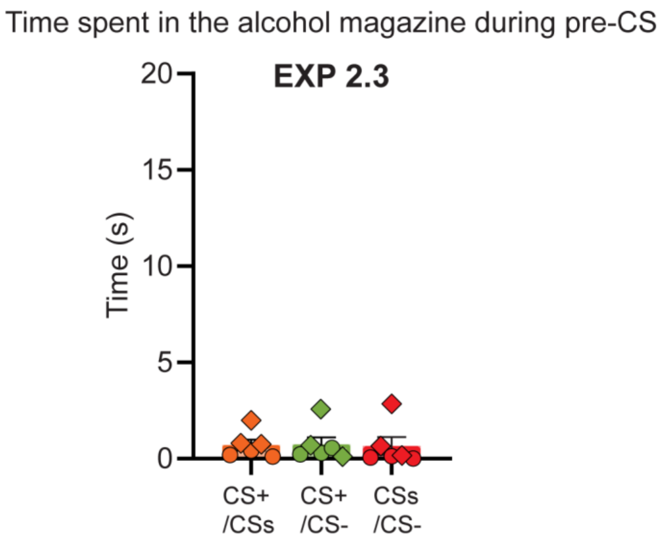


**Figure S5. Time spent in the alcohol magazine during pre-CS period in Exp. 2.3.** Mean ± standard error of the mean (SEM) time spent in the alcohol magazine during pre-CS in the conflict phase of experiment 2. (n = 3 males; diamond shapes, n = 3 females, circle shapes).
