## Supplemental tables for "Dual Role of LH-GABA Neurons in Encoding Alcohol Reward and Aversive Memories"

| Experimental Phase | Figure | Statistical Comparison | Significance time window  (Seconds; 0 = Cues turned on) |
| --- | --- | --- | --- |
| Exp. 1.1 Reacquisition | 2D | BCI: First session, CS+ | 0.35🡪8.50 |
|  | 2D | BCI: First session, CS- | 0.50🡪9.9 |
|  | 2D | PT: First session (CS+ v CS-) | -3.77🡪-2.2; 1.94🡪2.8 |
|  | 2E | BCI: Last session, CS+ | 0.38🡪6.29 |
|  | 2E | BCI: Last session, CS- | 0.62🡪10.75 |
|  | 2E | PT: Last session (CS+ v CS-) | 0.5🡪2.64 |
|  | 2F | PT: CS+ (First v Last Cond. session) | n.s. |
|  | 2G | PT: CS- (First v Last Cond. session) | n.s. |
| Exp. 1.2  Conflict | 3D | BCI: First session, CS+A | 0.38🡪5.03 |
|  | 3D | BCI: First session, CS+A+S | 0.38🡪7.98; 10.25🡪15.22 |
|  | 3D | BCI: First session, CS- | 0.44🡪3.58; 5.72🡪6.41 |
|  | 3D | PT: First session (CS+A+S v CS+A) | 10.25🡪13.96 |
|  | 3D | PT: First session (CS- v CS+A) | 0.57🡪3.9 |
|  | 3D | PT: First session (CS- v CS+A+S) | 1.01🡪4.34; 7.42🡪8.30; 10.25🡪15.34 |
|  | 3E | BCI: Last session, CS+A | 0.25🡪12.38; 13.45🡪20 |
|  | 3E | BCI: Last session, CS+A+S | 0.31🡪20 |
|  | 3E | BCI: Last session, CS- | 0.38🡪10.75 |
|  | 3E | PT: First session (CS+A+S v CS+A) | 10.31🡪13.5 |
|  | 3E | PT: Last session (CS- v CS+A) | 0.31🡪12.38; 13.89🡪15.34; 16.72🡪17.35; 17.48🡪17.86; 18.23🡪20 |
|  | 3E | PT: Last session (CS- v CS+A+S) | 0.38🡪16.10; 17.92🡪18.74 |
|  | 3F | PT: CS+A (First v Last) | 0.38🡪3.58; 4.53🡪6.47; 7.42🡪10.88; 14.2🡪15.22; 18.30🡪19.93 |
|  | 3G | PT: CS+A+S (First v Last) | 0.44🡪1.95; 4.4🡪6.98 |
|  | 3H | PT: CS- (First v Last) | n.s. |

**Table 1.** **Time periods (in seconds) where the photometry statistical tests are significant in Experiment 1.** *Time 0 refers to the start of the 20 second Cue Period. BCI, Bootstrapped Confidence Intervals; PT = Permutation Tests.*

| Figure | Comparison | Statistic | Adjusted p-value |
| --- | --- | --- | --- |
| 5B | CSa Pre Early vs. Late | t(5) = 2.42 | > 0.05 |
| 5B | CSa During Early vs. Late | t(5) = 5.47 | = 0.0028 |
| 5B | CSa After Early vs. Late | t(5) = 14.5 | < 0.0001 |
| 6B | CSa during vs. CS- during | t(5) = 2.32 | = 0.2 |
| 6B | CSa during vs. CSs during | t(5) = 3.15 | = 0.07 |
| 6B | CS- during vs. CSs during | t(5) = 1.63 | = 0.49 |
| 6C | CSa after vs. CS- after | t(5) = 12.7 | < 0.001 |
| 6C | CSa after vs. CSs after | t(5) = 14.13 | < 0.001 |
| 6C | CS- after vs. CSs after | t(5) = 2.43 | = 0.28 |
| 7B | Conflict vs. Reward during | t(5) = 2.86 | = 0.10 |
| 7B | Conflict vs. Shock during | t(5) = 1.21 | = 0.84 |
| 7B | Reward vs. Shock during | t(5) = 2.90 | = 0.10 |
| 7C | Conflict vs. Reward after | t(5) = 0.32 | = 0.99 |
| 7C | Conflict vs. Shock after | t(5) = 5.91 | < 0.01 |
| 7C | Reward vs. Shock after | t(5) = 4.87 | < 0.05 |

**Table 2. Post-hoc analyses of magazine entries during the different phases of experiment 2.**

| Experimental Phase | Figure | Statistical Comparison | Significance time window  (Seconds; 0 = Cues turned on) |
| --- | --- | --- | --- |
| Exp. 2.1 Reward only | 5D | BCI: CSa (Early) | 0.25🡪23.14; 27.98🡪29.12; 32.2🡪40 |
|  | 5D | BCI: CSa (Late) | Above baseline: 0.31🡪10.25; 15.09🡪15.53  Below baseline: 24.15🡪31.44 |
|  | 5D | PT: (Early v Late) | 0.25🡪4.0; 16.5🡪37.29 |
| Exp. 2.2 Discrimination | 6D | BCI: Alcohol (CSa) | Above baseline: 0.25🡪18.86  Below baseline: 23.46🡪37.23 |
|  | 6D | BCI: Control (CS-) | 0.25🡪40 |
|  | 6D | BCI: Shock (CSs) | 0.25🡪40 |
|  | 6D | PT: (Alcohol v Control) | 17.23🡪40 |
|  | 6D | PT: (Alcohol v Shock) | 0.31🡪14.21; 16.16🡪40 |
|  | 6D | PT: (Shock v Control) | 0.31🡪6.6; 10.44🡪11.63  19.75🡪22.08 |
| Exp. 2.3  Conflict | 7D | BCI: Conflict | 0.25🡪23.59; 37.11🡪40 |
|  | 7D | BCI: Reward | 0.25🡪20.81; 33.27🡪40 |
|  | 7D | BCI: Shock | 0.25🡪40 |
|  | 7D | PT: (Conflict v Reward) | 0.38🡪1.19; 2.64🡪10.06;  20.19🡪24.59; 32.83🡪33.96 |
|  | 7D | PT: (Conflict v Shock) | 12.14🡪13.21; 20.19🡪22.01; 24.49🡪34.97 |
|  | 7D | PT: (Shock v Reward) | 0.75🡪1.57; 18.93🡪33.15 |

**Table 3.** **Time periods (in seconds) where the photometry statistical tests are significant in Experiment 2.** *Time 0 refers to the start of the 20 second Cue Period. BCI, Bootstrapped Confidence Intervals; PT = Permutation Tests.*
